# ECM degradation in the stump region induced by Fgf during functional joint regeneration in frogs

**DOI:** 10.1101/2023.08.04.551975

**Authors:** Haruka Matsubara, Takeshi Inoue, Kiyokazu Agata

## Abstract

Previous studies indicated the importance of the reciprocal interactions between residual tissues in the stump and the newly formed blastema for achieving functional joint regeneration after amputation at the joint level in newts. This reciprocal interaction during regeneration was named “reintegration”. When this reintegration mechanism was evoked in *Xenopus leavis*, regeneration of a functional joint was induced in frogs. Interestingly, degradation of extracellular matrix (ECM) in the remaining joint cartilage was observed during regeneration in both newts and frogs. Histological and gene expression analyses suggested that the degradation of Type II collagen in the cartilage of the articular head might be performed by matrix metalloproteases (Mmps) which were transiently expressed after amputation. We found that fibroblast growth factor (Fgf) induced Mmps expression in the cartilage of the articular head. These results support the possibility that the Fgf signal induces ECM degradation in joint tissues via Mmps expression and that the ECM degradation and subsequent bone morphogenetic protein (Bmp) secretion promote cell proliferation, migration, and differentiation of the cells in the blastema to achieve functional joint regeneration.

## Introduction

Amphibians have remarkable regeneration capacities in various tissues and organs, such as the eyes, brain, heart, jaw and limbs (Inoue et al., 2012; Giudice et al., 2008; Kurosaka et al., 2008; Urata et al., 2018; Joven et al., 2019; Uemasu et al., 2022; Ikuta et al., 2023), although the regeneration capacities of the limb differ between species. Urodele amphibians, namely, newts and salamanders, can regenerate the complete limb structure when a limb is cut in any region. The regeneration capacities of urodeles are sustained after metamorphosis (Agata & Inoue 2012; Inoue et al., 2012). On the other hand, anuran amphibians have decreased regeneration capacity after metamorphosis (Phipps et al., 2020). For example, the African clawed frog, *Xenopus laevis*, can regenerate almost the complete hindlimb structure when it is cut until metamorphoses proceeding, but after metamorphosis, their regeneration ability decreases, and they cannot regenerate the proper limb structure, and instead produce a cartilage rod called a ‘spike’, regardless of whether the lower arm or upper arm is cut (Dent, 1962; Endo et al., 2000; Hayashi et al., 2015). The spike does not contain any morphologically or functionally normal structures, including joints (Tassava, 2004; Satoh et al., 2005).

Despite this difference of limb regeneration capacities between urodele and frog, several signaling molecules were commonly involved in the induction and maintenance of the blastema, a mass of dedifferentiated cells (Han et al., 2001; Tsai et al., 2020). Fibroblast growth factor (Fgf) is one of the most important signaling molecules for forming the blastema and is produced by nerves and wound epithelium/apical epidermal cap (AEC) (Mullen et al., 1996; Maddaluno et al., 2017). The AEC is a regeneration-specific structure that corresponds to the apical epidermal ridge (AER) observed in the developing limb (Christensen and Tassava, 2000; Satoh et al., 2012; C. McCusker et al., 2015). In the AEC, several Fgfs are expressed and regulate cell proliferation and dedifferentiation in the regenerating stump in amphibians, including frogs (Christen and Slack, 1997; Endo et al., 2000; Yokoyama et al., 2000; Cannata et al., 2001; Han et al., 2001; Okumura et al., 2019; Aztekin et al., 2021;). Several studies indicated that limited gene expressions or cellular potencies cause spike regeneration in frogs (Endo et al., 2000; Yakushiji et al., 2007 and 2008; Ohgo et al., 2010; Aztekin et al., 2021; Lin et al., 2021).

Also, it had been thought that frogs could not regenerate any joint structure in the spike after metamorphosis (Tassava, 2004; Satoh et al., 2005). However, our previous study demonstrated that *Xenopus laevis* had the potential to regenerate functional joints (Tsutsumi et al., 2016). When the froglet forelimb was amputated at the elbow joint level, the remaining and the regenerating cartilage made proper connection to remake the functional joint. In addition, muscles and tendons from the remaining part could make proper connections to the regenerated cartilage. During the functional joint regeneration in the urodele limbs, the proximal residue (named hereafter the “stump”) affects and interacts with the blastema which was newly formed between stump and AEC: this was named “reintegration” (Tsutsumi et al., 2015). The joint tissues contain several types of tissues (muscle, cartilage, ligament, and tendon) and these tissues’ orchestration is necessary for “functional” joint regeneration. Proper tissue connections between the stump and the blastema might be achieved by the reintegration mechanisms. However, the stump-to-blastema interaction has not been investigated over a long period, although the reintegration mechanisms may work in all regeneration processes. Thus, here we approached the molecular mechanisms of the reintegration which drive the interaction between the stump and the blastema. Especially, we focused on the cellular dynamics and signaling networks in the stump during joint regeneration since we found ECM degradation in the remaining articular cartilage during functional regeneration (Tsutsumi et al., 2015 and 2016). We found that, during joint regeneration in *Xenopus laevis*, the remaining stump tissues might be activated by signals derived from the blastema to produce signaling molecule(s) which actively regulate cellular fates in the blastema to achieve proper tissue-tissue connection for functional joint regeneration.

## Materials and methods

### Animals

*Xenopus laevis* froglets were purchased from Watanabe farm or raised from fertilized eggs (a kind gift from Harumasa Okamoto). *Xenopus laevis* froglets for blastema transplantation were wild-type (WT) (J-strain) and transgenic (JG-hybrid (J/GFP (green fluorescent protein))), provided by Hitoshi Yokoyama, Hirosaki University and National BioResource Project (NBRP, Clawed frogs / Newts) Hiroshima University Amphibian Research Center. The animals were kept in plastic containers at room temperature (25 °C) and fed daily. The animals were handled with an animal care protocol of Gakushuin University and the Faculty of Medicine, Tottori University.

### Limb amputation and sampling

The animals were anesthetized with 0.2% ethyl 3-aminobenzoate methanesulfonate salt (MS-222) (Sigma-Aldrich) for the limb amputation and sampling at various times thereafter. Joint amputation was described previously (Tsutsumi et al., 2016). After the limb was cut, the animals were fed daily until the time of sampling. Photographs were taken using a Leica M125 (Leica Microsystems, Wetzlar, Germany). Ethical treatment of experimental animals was conformed to the policy on the care and use of laboratory animals of Gakushuin University and the Faculty of Medicine, Tottori University.

### Blastema transplantation

To distinguish between the donor and host cells, we used J-strain WT (donor) and J/G-hybrid (J/GFP) (host) in the experiment to avoid immune rejection (Otsuka-Yamaguchi et al., 2017). The animals were allowed to regenerate until 12-14 days post amputation (dpa), and then the blastema was dissected and the blastema was explanted at the stump region, The remaining explanted blastema could be confirmed by the GFP fluorescence. Transplanted animals were heat-shocked (34°C, 30 min) 6 h prior to fixation. Samples were harvested and sectioned 12-14 days post transplantation.

### Frozen section preparation

Samples were fixed with 4% paraformaldehyde/phosphate-buffered saline (PBS) at 4 °C overnight and decalcified for several days in G-Chelate Mild (GenoStaff). After decalcification, samples were soaked serially in 20% and 30% sucrose/PBS, and finally embedded and frozen in O.C.T. compound (Sakura Fine Technical Co. Ltd., Japan). Frozen samples were sectioned using a cryostat (Microm HM550; Thermo Scientific and CM1950; Leica) at 10 μm thickness.

### Histological staining

Elastica van Gieson (EVG) staining was performed on frozen section samples. Sections were post-fixed using 4% paraformaldehyde /PBS before staining.

### Immunofluorescence staining

Frozen section samples were rinsed with PBS three times for 5 min each and blocked with 3% bovine serum albumin (BSA)/PBS for 1 h. For type I and type II collagen staining, samples were incubated with 10 μg/ml Proteinase K/PBS at 37 °C for 30 min to induce antigen retrieval before blocking. After blocking, sections were incubated with primary antibody diluted in 3% BSA/PBS, at 4 °C overnight. Primary antibodies were mouse anti-collagen type I monoclonal antibody (Sigma-Aldrich, #C 2456, diluted 1:100), mouse anti-collagen type II monoclonal antibody (Developmental Studies Hybridoma Bank, #II-II6B3, diluted 1:100), rabbit anti-laminin polyclonal antibody (Sigma-Aldrich, #L9393, diluted 1:500), mouse anti-myosin heavy chain monoclonal antibody (Developmental Studies Hybridoma Bank, #MF20, diluted 1:100), mouse anti-GFP monoclonal antibody (Fujifilm Wako, mFX73, diluted 1:100), Rat anti-GFP monoclonal antibody (nacalai tesque, GF090R, diluted 1:1000) and rabbit anti-pSmad1/5/9 antibody (Cell Signaling Technology, #13820, diluted 1:100). After the primary antibody incubation, sections were incubated with Alexa Fluor 488-, or 594- (Invitrogen) or HRP- (Cell Signaling Technology, #7074) labeled secondary antibodies (diluted 1:100). In the case of pSmad1/5/9 staining, PhosSTOP (Roche) and Tyramide Signal Amplification Kits (Invitrogen) were used.

### *In situ* hybridization

Probe sequences for *in situ* hybridization with *Xenopus laevis* gene transcripts were isolated by PCR using cDNA generated from *Xenopus laevis* regenerate blastema (15 dpa). The primer sequences used were *mmp1* (forward primer, CAG ATG TGG AGT CTA TGA TGT TGG; reverse primer, GAA ATC GTT GCT TAT TTG CTT TGG; NM_001094954.1), *col1a2* (forward primer, GAA ACT TTG CTG CTC AGT ATG ACT C; reverse primer, CCA CTA CGA CCC TCA ATA CCT TG; NM_001087258.1), *col2a1* (forward primer, GCA TGA AGA CAA GAT AAC TAT TGG AGA C; reverse primer, AGG TGG TCC TCT TGG TCC TAA AAC; NM_001087789.1), *prg4* (forward primer, GGT GGA GGA TAGT ACA GAC GAT C; reverse primer, CTT ATA TGC TGG AGG TCA CTC TCT GAT A; XM_018259871.1), *prrx1*(*prx1*) (forward primer, AAG AGT GTT TGA GAG GAC GCA TTA TC; reverse primer, AAT TGA CTG TTG GCA CTT GGT TCC; [Takahashi et al., 1998] XM_018258780.1)

*In situ* hybridization was performed as previously reported (Tsutsumi et al., 2016; Matsubara et al., 2017). Signals were detected using an inverted routine microscope ECLIPSE Ts2-FL (Nikon), an Imager M1 microscope and Axiocam MRc5 camera (ZEISS), BZ-9000(KEYENCE).

### Quantitative RT-PCR (QRT-PCR)

The distal blastema and proximal stump region of experimental samples were separated from each other. Control samples were obtained from the forelimb by cutting at the midpoint of the humerus. Each type of sample was prepared in triplicate. RNAs were isolated using an RNAeasy Micro Kit (Qiagen). To homogenize samples, we used a Minilys homogenizer (Bertin Instruments) and a beads kit (CK14; Bertin Instruments) under the conditions of 5000 rpm, 30 sec, 2∼3 times. After RNA isolation, we prepared cDNA using a QuantiTect Reverse Transcription Kit (Qiagen). Quantitative RT-PCR was performed using an ABI Prism 7900HT PCR Sequence Detection System (Applied Biosystems) or ViiA7 (Thermo Fisher Scientific). Getting data was statistical analysised and calculated using the ddCT method by normalization to *gapdh* expression. The primers were *mmp1* (forward primer, ACC AGA ATC TAC AAT GAA GTA TCA GAT ATA; reverse primer, GAA ACA GAT TGT ATA TTT CAC TGG TCT TTG; NM_001094954.1), *mmp9* (forward primer, AAG ACA TTG ACT ATT GAT AAT GGC TAC C; reverse primer, AGT TTC CTT GAT AGA GAA AAA CAT CAT G; NM_001086503.1), *mmp13* (forward primer, GAA GAT GAA AGA TGG ACA AGT TCT AGT G; reverse primer, GTT CAC ATA ACG ATA GTT TGG AAA CAT C; NM_001086405.1), *timp1* (forward primer, TGC AGA TTT TGT TAT TCG AGG GAG ATT C; reverse primer, ATC TCA TAT TTC ACT CTG CCC TGC TCT T; NM_001287451.1), *col1a2* (forward primer, AAT TAC CCG TTA AGA AGT GTA TGA AAG C; reverse primer, TTA CAG ATT GGC ATG TTG CTA GGT ATA G; NM_001087258.1), *col2a1* (forward primer, GCA TGA AGA CAA GAT AAC TAT TGG AGA C; reverse primer, GAA TCC ACA AAG CTA AAC ATG TCT AAA T; NM_001087789.1), *prg4* (forward primer, CTA CTG TGA AGG CCA TTT CTG TTG; reverse primer, CTG GAG GTC ACT CTC TCT GAT ATC CC; XM_018259871.1), *prrx1* (forward primer, AAG AGT GTT TGA GAG GAC GCA TTA TC; reverse primer, CTC TGG CTT CAG TGA GGT TAA CTC TG; XM_018258780.1), *fgf2* (forward primer, AGT GAC TTG CAC ATA AAA TTA CAG CTA C; reverse primer, AAG AAG CAT TCA TCT GTT ATA CAC TTC A; NM_001099871.1, XM_018243334.2), *dusp6* (forward primer, ATA TTG AAT CTG ACA TTG ACA GGG ATC; reverse primer, CCA GGT TAG TAG AAT CCT TAG CAC AAC; NM_001094761.2), *gapdh* (forward primer, GAC CAG GTT GTC TCC ACT GAC TTC A; reverse primer, CCG CAT TCA TTA TCA TAC CAG GAA; NM_001087299.1).

### EdU cell proliferation assay

Animals were injected intraperitoneally with 100 μl of 1 mM EdU (5-ethynyl-2’-deoxyuridine) 2 days before sampling. Samples were processed for frozen sections and EdU incorporation was detected by using a Click-iT EdU imaging kit (Invitrogen).

### Beads implantation in the joint capsule

Affi-Gel Blue Gel beads (Bio-Rad) were incubated with 1 mg/ml Fgf2 (recombinant human basic Fgf) (Fujifilm Wako) or PBS for at least 1 hour at 4 °C. To approach the elbow joint capsule, the dermis was cut near the elbow with micro spring scissors from the ventral region and then a notch was cut in the joint capsule with a surgical knife. The beads were implanted using an electropolished tungsten needle as described by Yu et al., 2010.

## Results

### Changes of gene expression in the stump during joint regeneration

In joint regeneration, previously we found residual articular cartilage ECM degradation in the stump region, both of newt and frog (Tsutsumi et al., 2015 and 2016). To reveal the details of gene expression changes of the stump tissue during joint regeneration, we performed QRT-PCR at 3 time points of joint regeneration, namely, wound-healing stage (5 days post amputation (dpa)), early-regenerate stage (10-11 dpa), and middle-regenerate stage (15-16 dpa) (Fig. 1). Tissue structural changes in each time point were checked by Elastica van Gieson (EVG) staining and COL1/Laminin immunostaining (Supplement 1-1). We screened several regeneration-related genes in the stump represented by using tissue markers, ECM degradation enzymes, signaling molecules, and signal-related genes, and comparing these gene expression levels in the joint stump to those in the normal (uncut) forelimb. Several genes showed upregulation in the joint regeneration stump. Several tissue degradation enzyme matrix metalloproteinase (Mmp) genes were upregulated in the joint stump. The expression levels of Mmp genes *mmp1*, *mmp9*, *mmp13*, and *timp1* (tissue inhibitor of metalloproteinase 1) were slightly upregulated in the wound-healing stage (Fig. 1). In the early-regenerate stage, Mmp genes were upregulated to the maximum observed levels. In the middle-regenerate stage, which is the stage of the completion of ECM degradation, Mmp gene expression levels returned to the wound-healing stage levels (Fig. 1). These findings indicate that Mmps are among the tissue degradation enzymes in the joint stump. Moreover, several tissue-specific (*col1a2*; tendon and ligament, *prg4*; synovium membrane) and regenerate blastema-specific (*prrx1*) genes were upregulated in the early-regenerate stage of the joint regenerate stump (Fig. 1). The timing of the upregulations of these tissue-specific cells and blastema marker genes corresponded to the timing of the maximal Mmp genes upregulation. These findings suggest that tissue degradation in the stump may be the trigger for the tissue cells’ dedifferentiation and blastema formation in joint regeneration.

**Fig. 1.**
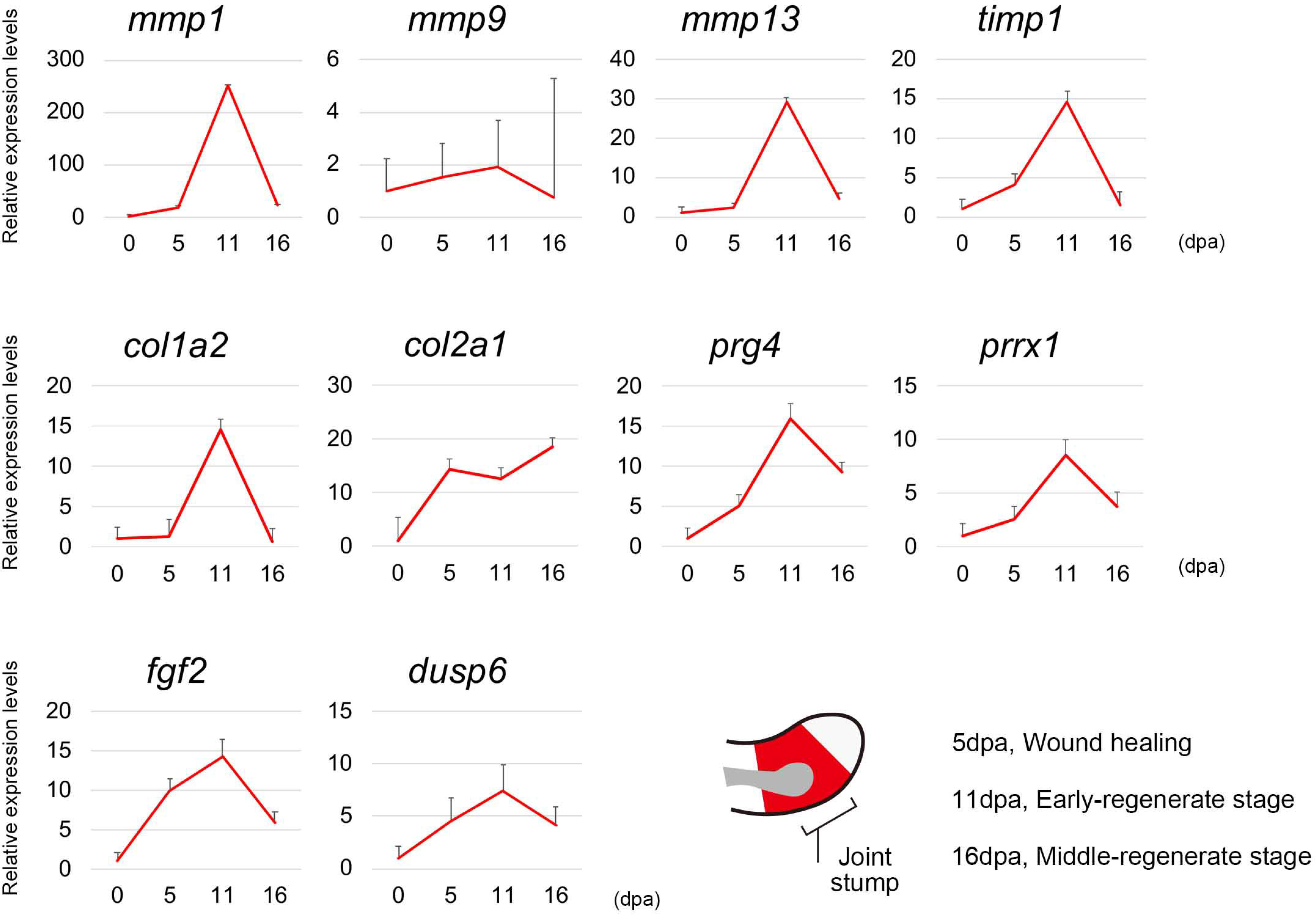
Genes expression in the stump of joint regeneration. Each graph indicates relative expression patterns of the intact forelimb sample (distal to the half of the humerus, 0 days post amputation (dpa)). Samples were taken at each indicated time point (3 sample mix, n=3). Bars: S.E.

### Mmp and ECM genes show joint regeneration-specific expression patterns, especially in the stump region

Next, to reveal the relationship between tissue structure changes and highly regenerative responses, we performed *in situ* hybridization for each upregulated gene. *mmp1* expression was detected in the joints’ articular head during the wound healing (Fig. 2A). The number of the *mmp1*-expressing cells in articular head was increased in the early-regenerate stage (Fig. 2B). In the middle-regenerate stage, the expression level of *mmp1* was drastically decreased to an undetectable level (Fig. 2C). These changing patterns of expression levels were consistent with the QRT-PCR analysis (Fig. 1). The *mmp1*-expressing region corresponded to the region of Col2 degradation (Fig. 2E asterisk), suggesting that the degradation of Col2 in the articular cartilage was caused by *mmp1* in a cell-autonomous manner. In the normal spike, lower arm regeneration, faint *mmp1* expression was detectable in the blastema (Supplement 2-1A, A’). Then we analyzed the expression of *col1a2*, *col2a1*, *prg4*, and *prrx1* genes in the joint of the early regenerate stage. The connective tissue cell marker *col1a2* was strongly expressed in the reforming joint capsule, including the blastema (Fig. 3A). The highly *col1a2*-expressing regions extended to the stump. Moreover, these cells from the stump were positive for expression of *prrx1*, a blastema cell marker (Fig. 3D, asterisk). These findings may indicate that stump-derived dedifferentiated cells actively contribute to the blastema and would be the main source of the regenerate joint capsule. In the normal spike, strong *col1a2* expression can detect from the stump to the blastema, although *prrx1* expression cannot detect the stump connective tissues except for the periosteum (Suzuki et al., 2005) (Supplement 3-1A-B’). The cartilage cell marker *col2a1* was expressed in the stump region of the articular cartilage (Fig. 3B). *col2a1* was expressed constantly in normal articular cartilage in the elbow joint (Supplement 3-1C). This *col2a1* homeostatic expression in the joint articular cartilage may reflect high cartilage-reforming ability after degradation by Mmps during joint regeneration. The synovial membrane marker gene *prg4* was expressed in the most basal layer of the reforming joint capsule (Fig. 3C). This indicates that synovial membranes were already regenerated (or repaired) before the other joint structures regenerated. These specific responses indicate that stump-to-blastema wide responses and remaking of joint tissue occurred in joint regeneration.

**Fig. 2.**
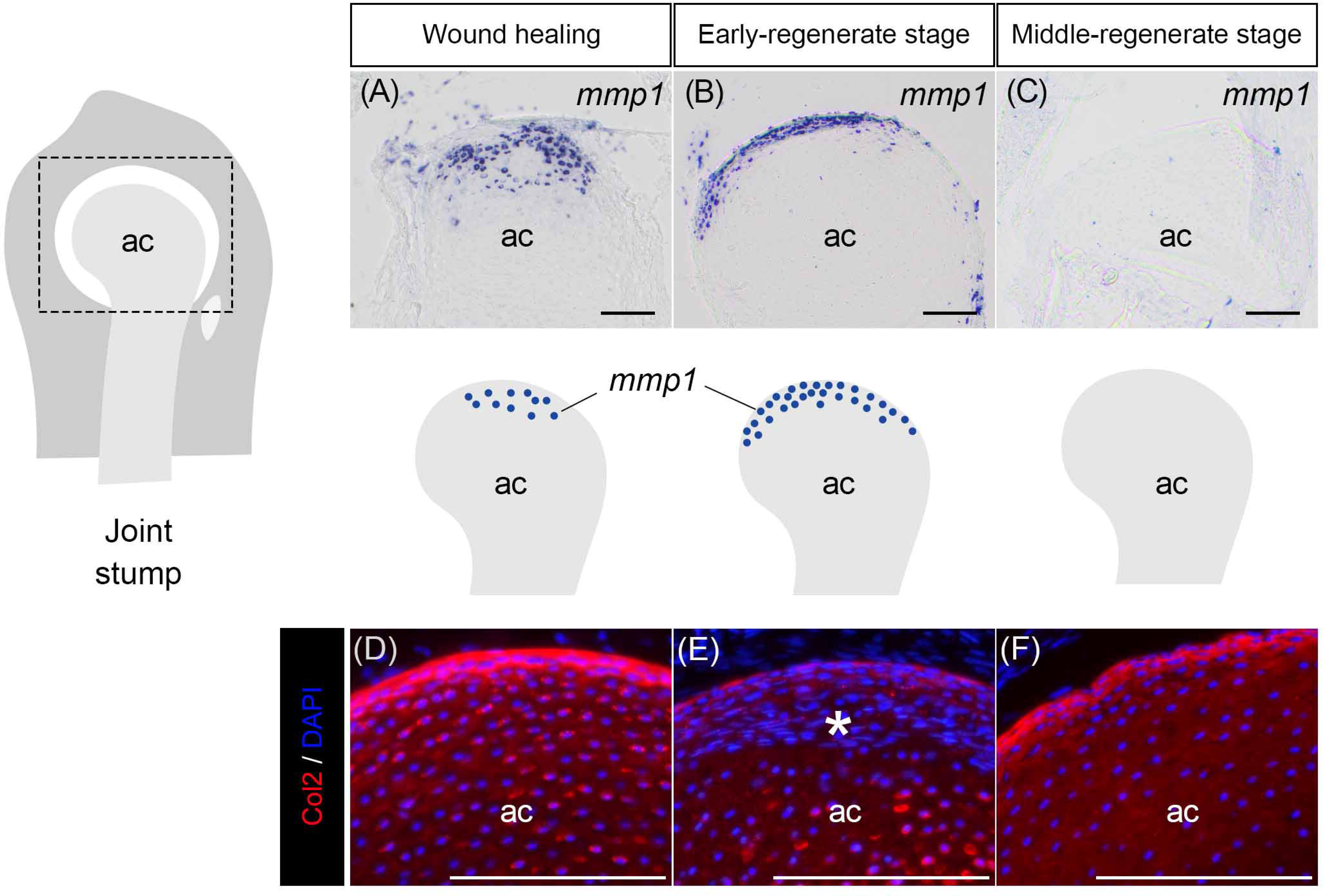
*mmp1* and Col2 expression patterns during joint regeneration in the stump. (A-C) The section *in situ* hybridization to detect *mmp1* expression and schematic diagram of the wound healing until the middle-regenerate stage for articular cartilage during joint regeneration. (D-F) Col2 expression in the peripheral region of articular cartilage during joint regeneration. (E) The asterisk indicates a low Col2-staining region in articular cartilage. ac, articular cartilage. Scale bars, 100 μm.

**Fig. 3.**
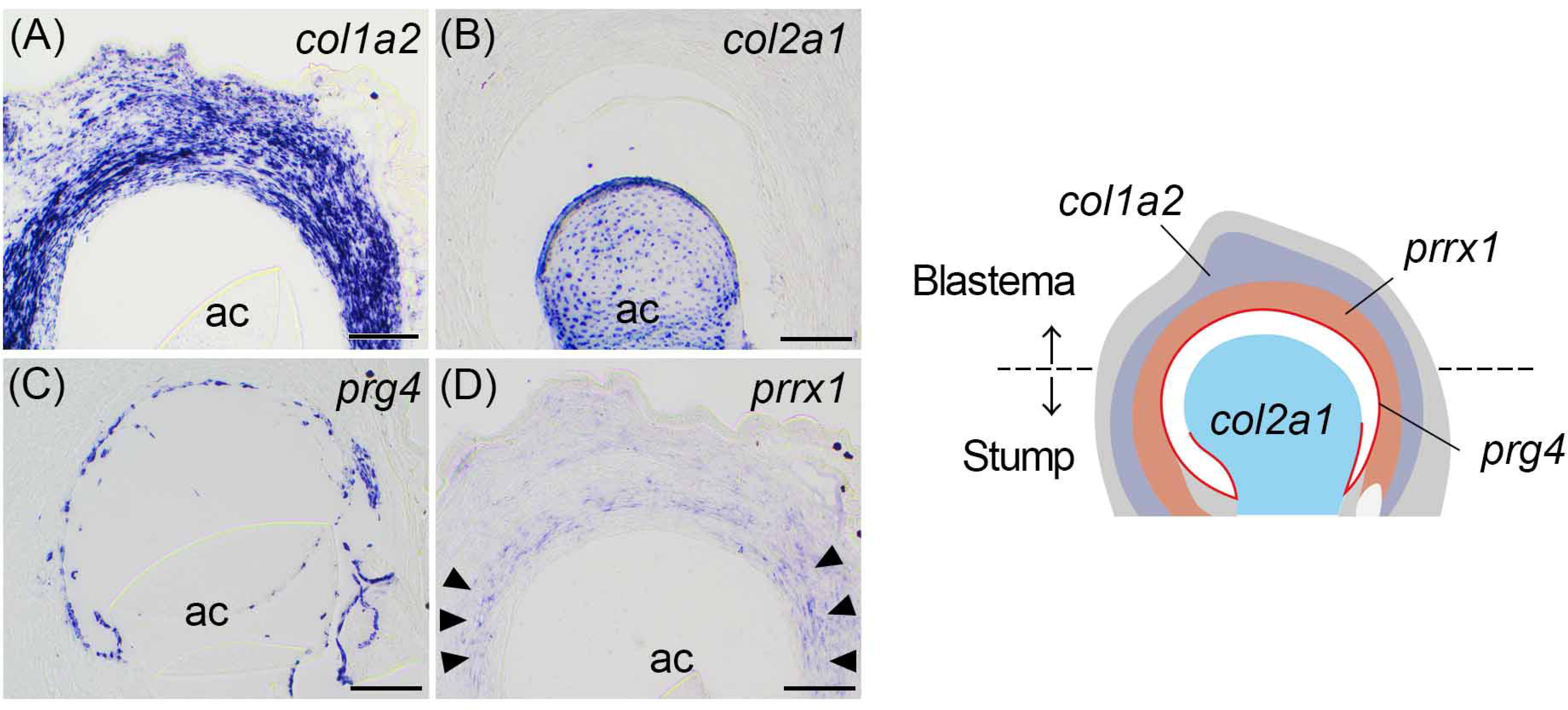
Gene expression patterns of joint regeneration. (A-D) The section *in situ* hybridization of the *col1a2*, *col2a1*, *prg4*, *prrx1* expression in the early-regenerate stage stump to blastema region of joint regeneration. Schematic diagram of the gene expression patterns in the early-regenerate stage of joint regeneration. Asterisks indicate the stump expression of *prrx1*.ac, articular cartilage. Scale bars, 100 μm.

### Active cell proliferation in the stump region during joint regeneration

To reveal specific cellular behavior in the stump of joint regeneration we compared cell proliferation in the stump of the joint and lower arm regeneration (Fig. 4A). Here, we used the criterion of 500 µm distance from the distal tip of the blastema to discriminate the stump in each type of regeneration. The results showed that two-fold higher cell proliferation was detected in the stump region of joint regeneration compared to lower arm regeneration (Fig. 4B). One of the reasons for the difference in the rate of cell proliferation between them was that the composition of tissues in these stumps is different. The joint stump region has abundant connective tissues such as tendons and ligaments. On the other hand, the lower arm stump is mainly comprised of only muscles and skeletal callus (Miura et al., 2015). The result indicates that the cells in the stump of the joint actively proliferate, which may be one of the reasons for the high contribution of the stump to the blastema in the joint regeneration.

**Fig. 4.**
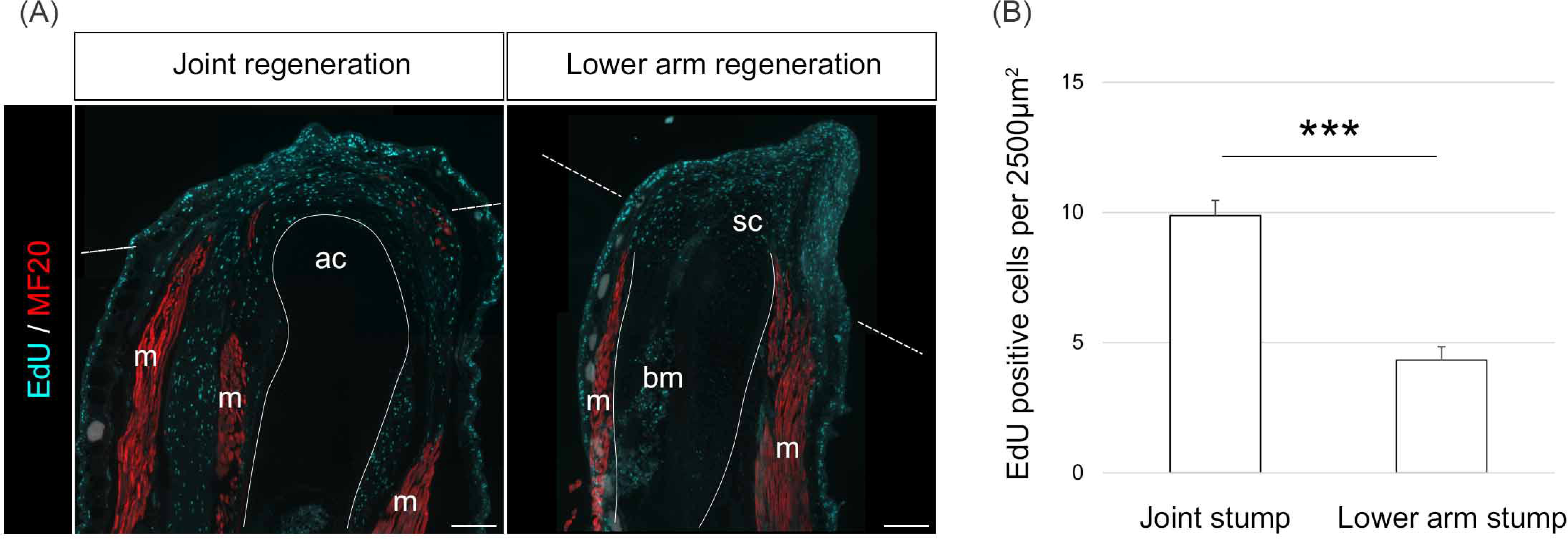
EdU assay of joint regeneration. (A) Cell proliferation detect by the EdU incorporation (Cyan) stained with muscle marker MF20 (Red) in joint and lower arm regeneration in the early-regenerate stage (10-11 dpa). The dashed line indicates the junction between blastema and stump. ac, articular cartilage; m, muscle; sc, spike cartilage; bm, bone marrow. (B) EdU-positive cell number in areas of 2500µm^2^ in the stump (within 500 µm from the distal tip) regions. Bars: S.E. ***P<0.01. Scale bars, 100 μm

### The remaining stump is crucial for the functional joint regeneration

To reveal the amount and period of contribution from the stump to the blastema in the reintegration process of joint regeneration, we performed a blastema transplant experiment of joint regeneration. We transplanted the blastema of the wild-type (WT) joint amputation to the stump of the JG-hybrid (J/GFP) joint amputation (Fig. 5A). The 12 dpa blastema was used as the transplant. Transplanted blastema remained and contributed to the regeneration 14 days after transplantation (Fig. 5B). To further reveal the cellular contribution of the stump during the regeneration, we made slide sections of the tissues (Fig. 5C-D’). J/GFP was used to label cells in the stump side. Abundant GFP-positive cells derived from the stump were detected between the blastema and the stump region (Fig. 5D, D’). In this experiment, at the time of transplantation, the host proximal element contained GFP-positive articular cartilage, muscles, tendons, and ligaments. From this view, the host’s stump-derived GFP-positive muscles and connective tissues (tendons and ligaments) passed over the junction between the host (stump) and the donor (blastema), and contributed to the blastema region. These data suggest that the stump contributes to the distal region even after the blastema formation during the functional joint regeneration. The cells derived from the joint-stump tissues make an active cellular contribution, and the stump plays a key role in producing functional regeneration in the joint.

**Fig. 5.**
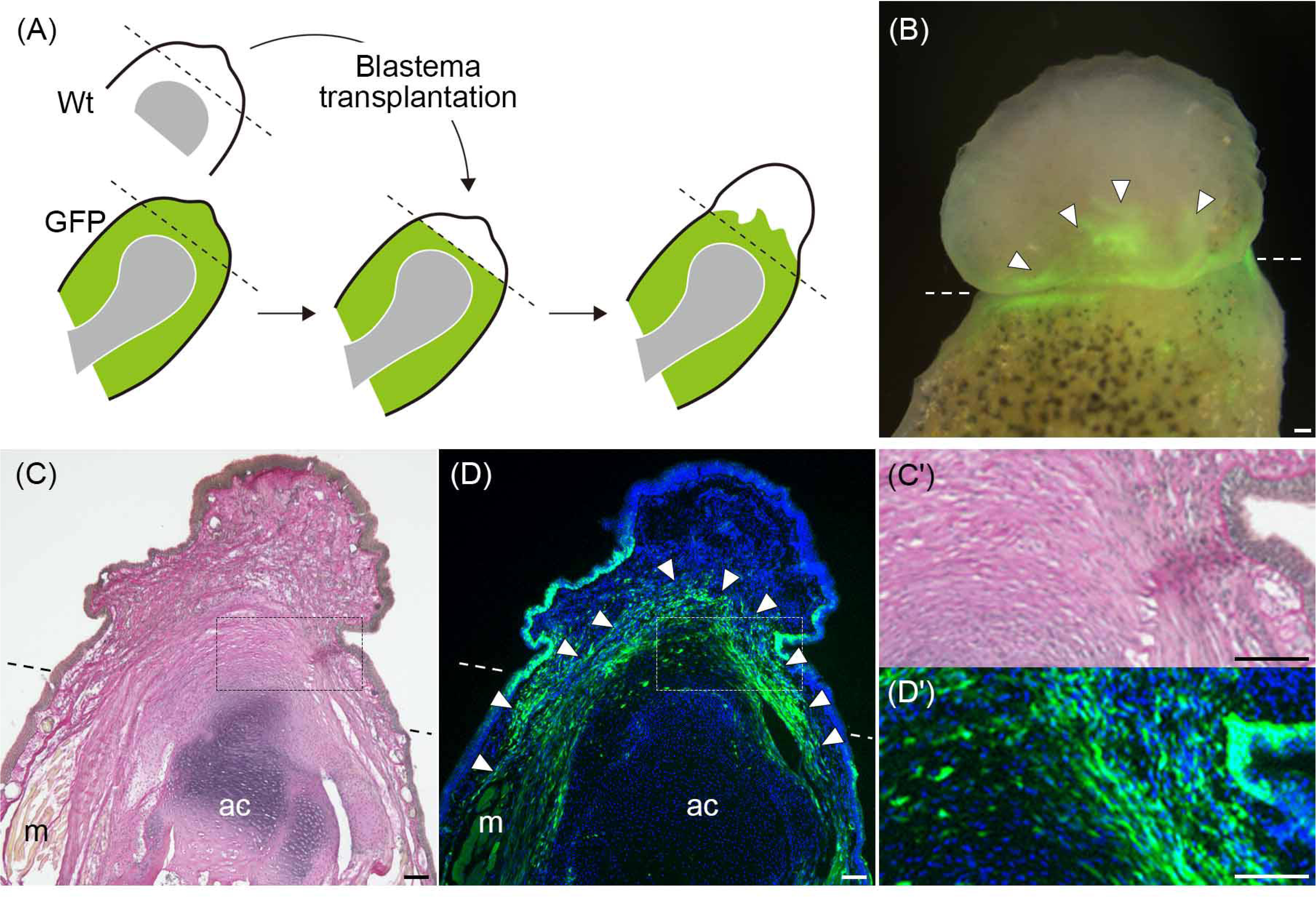
Blastema transplantation of joint regeneration. (A) Schematic diagram of the experiments. Each blastema was dissected and transplantation was performed. After transplantation, investigate the GFP cell’s contribution. (B) 14 days post transplantation (26 dpa) sample. The donor (blastema) was GFP negative, and the host (stump) was GFP positive. (C-D’) Section of the transplant experiment. (C, C’) EVG staining. (D, D’) GFP immunostaining and DAPI for nuclear staining. (C’, D’) Highly magnified view of the boxed area in (C, D). A dashed line indicates a junction of the blastema to the stump. Arrowheads indicate the GFP cell’s contribution to the blastema. ac, articular cartilage; m, muscle. Scale bars, 100 μm.

### Fgf can induce *mmp1* expression in the joint

Next, we examined what kind of molecular factor(s) induced joint regeneration activity. The gene expression analysis indicated that limb regeneration-signal-related genes were upregulated in the stump region of joint regeneration. Especially, Fgf signaling is a well-known limb regeneration inducer (Christen and Slack, 1997; Yokoyama et al., 2000; Cannata et al., 2001; Aztekin et al., 2021). Also, it was reported that MMPs expression can induced by Fgfs in the blastema of limb regeneration (Satoh et al., 2011). From these reports, we postulated that Fgf can induce Mmp-related gene expression during joint regeneration. Expression of *fgf2* and *dusp6* (a negative feedback regulator for Fgf signaling) was detected in the stump of joint regeneration (Fig. 1). To test the possibility of a relationship between Fgf signaling and Mmp-related tissue degradation, we implanted Fgf-soaked beads in the unamputated joint cavity of the elbow (Fig. 6A). *mmp1* expression was induced in the area proximal to the Fgf-soaked beads in the articular cartilage and synovial membrane, the same as in joint regeneration, although no *mmp1* expression was detected in the PBS-soaked bead implantation (Fig. 6B). Our QRT-PCR results indicated that upregulation of Fgf signal and MMPs expression were synchronized in the stump region of joint regeneration (Fig. 1). The results showed that the activation of Fgf signaling by joint amputation induces the upregulation of Mmp-related genes for articular cartilage degradation of the residual joint tissues. This indicates that regenerative signals reached the stump region of joint regeneration and tissue degradation could be induced in that region. This stump reaction and gene-activation relay could induce subsequent high regeneration potency in joint regeneration.

**Fig. 6.**
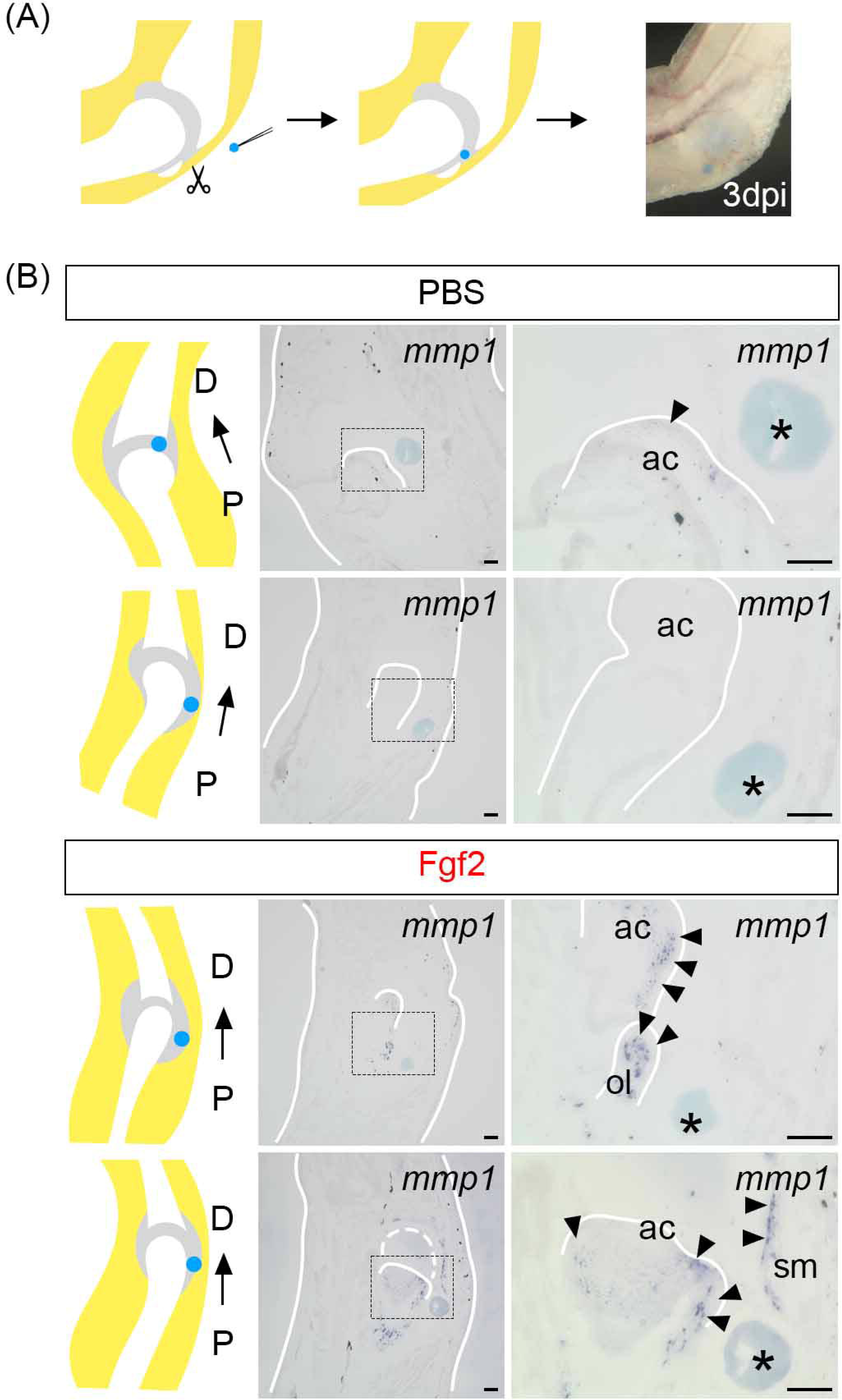
Fgf can induce *mmp1* expression in joint. (A) Experimental scheme of beads implantation to the elbow joint capsule. dpi: days post implantation. (B) *mmp1* expression in control (PBS-soaked beads) and Fgf2-soaked beads implanted samples. Arrowheads indicate positive signal regions and asterisks indicate beads. D, distal; P, proximal; ac, articular cartilage; sm, synovial membrane; ol, olecranon. Scale bars, 100 μm.

### Bmp signaling might be activated in basal blastema region of joint regeneration

Finally, we tried to investigate the role of the articular cartilage degradation in the stump of joint regeneration. One of the possibilities was that ECM degradation can release some signaling factors (Hynes, 2009). If there is release of signaling factor(s), they may affect surrounding regenerating tissues and may evoke cellular response(s). It is known that cartilaginous ECM Col2 can bind to Bmp2 through the chordin-like domain and change the binding status depending on the ECM property (Zhu et al., 1999; Xu et al., 2017). Therefore, we tried to detect Bmp signaling activation in joint regeneration by immunostaining of phosphorylated Smad 1/5/9 (pSmad1/5/9). There was almost no nuclear pSmad1/5/9 signal in the wound-healing stage (Fig. 7A, A’). In the early-regenerate stage of joint regeneration, the timing when strong ECM degradation occurred in the articular cartilage, some cells in the blastema adjacent to the cartilage showed nuclear pSmad1/5/9 (Fig. 7B, B’). The number of nuclear pSmad1/5/9-positive cells in the blastema was increased in the middle-regenerate stage (Fig. 7C, C’). These results indicate that the timing of the start of Bmp signaling activation corresponded to the tissue degradation timing in joint regeneration. pSmad1/5/9 activation was detected only in the neighboring cells of degraded articular cartilage. This regional specificity of Bmp signal activation may indicate that the degraded articular cartilage is one of the sources of Bmp and that degraded articular cartilage can induce Bmp signal activation in the surrounding blastema cells. It is known that Bmp is necessary to make joint structures during limb development and regeneration (Francis-West et al., 1999; Satoh et al., 2005; Yu et al., 2019). This suggests the possibility that Bmps from the degraded stump cartilage can induce not only cartilaginous differentiation but also the joint structure regeneration in joint regeneration.

**Fig. 7.**
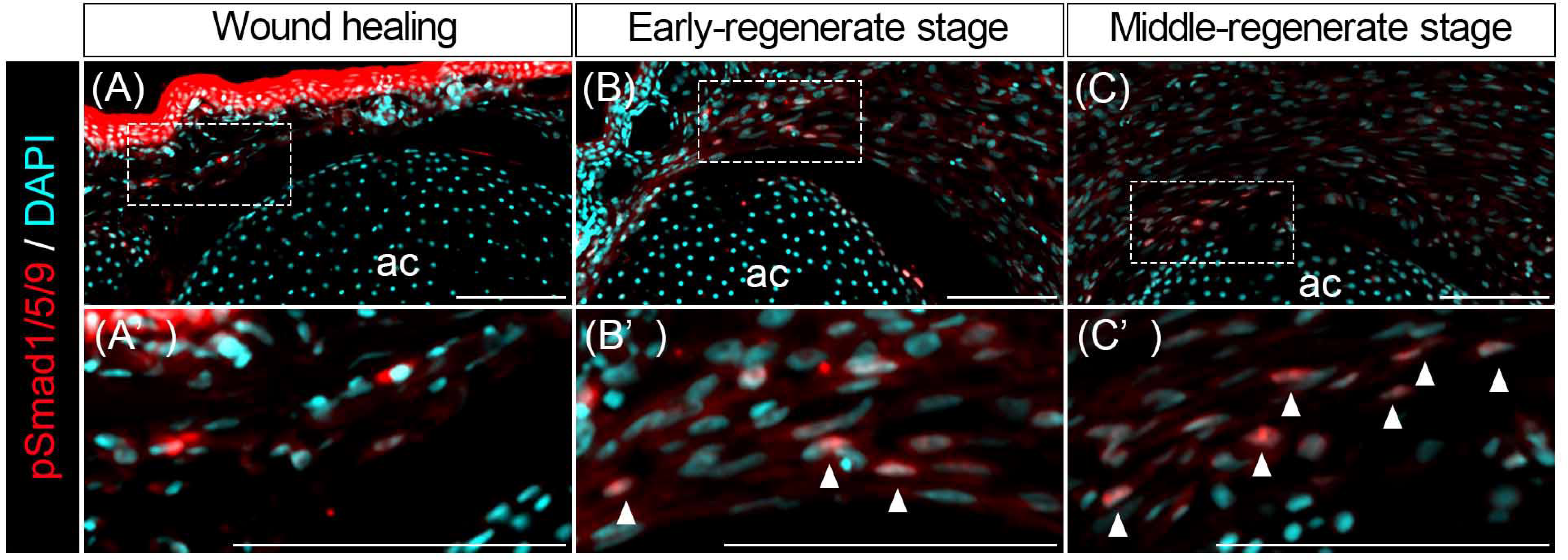
Bmp signal activation in joint regeneration. (A-C) Phosphorylated Smad1/5/9 (pSmad1/5/9) expression at each timing during joint regeneration. (A’-C’) Highly magnified view of the peripheral region of the articular cartilage. Arrowheads indicate nuclear accumulation of pSmad1/5/9-positive cells. ac: articular cartilage. Scale bars, 100 μm.

## Discussion

In this study, we revealed some tissue structure changes and signal molecule functions in the mechanism of joint regeneration by focusing on the stump tissue’s behaviors. In joint regeneration, stump tissues actively contributed to the blastema and responded to the regeneration signals. Furthermore, Fgf could induce *mmp1* expression in the stump articular cartilage.

### Stump responses in joint regeneration

In joint regeneration, tissue degradation enzyme *mmp1* expression was detected mainly in the articular cartilage of the stump (Fig. 2). Corresponding to the *mmp1* expression, Col2 degradation can detect in the articular head (Fig. 2E). These suggest *mmp1* in the stump articular cartilage should degrade Col2 (Fig. 2B, E). In limb regeneration, it is thought that the MMPs are the main player in ECM degradation (Yang et al., 1999; Vinarsky et al., 2005; Calvs et al., 2010). One possibility of MMP function in the stump of joint regeneration is dedifferentiation induction from the tissues. Upregulation of tissue stem cell-specific and regeneration-related genes in joint regeneration showed high activity in the stump (Fig. 1, Fig. 3). *Prrx1(Prx1)* is known to be expressed in the blastema (dedifferentiated cells) of lower arm regeneration (Suzuki et al., 2005 and 2007; Yokoyama et al., 2011). In joint regeneration, *prrx1* expression can be detected not only in the blastema but in the stump (Fig. 3D). These suggest joint stump shows regenerate responses even after the blastema formation.

### Stump contribution to the blastema in joint regeneration

There is no doubt that all blastema cells come from the amputated stump region. But in limb regeneration, it is not clearly known how long and how many stump tissues or cells contribute to regeneration. Remaking the proper size, shape, and functional integrity of structures is important for regeneration. Reintegration is an important aspect of regeneration, and it has been reported that the joint region is where reintegration can be clearly recognized (Tsutsumi et al., 2015 and 2016). This is because to make a functional joint, some tissues (such as muscles, tendons, and ligaments) or tissue-derived dedifferentiated cells in the stump are necessary to pass over the remaining segment and contribute to the blastema. In cell proliferation assays showed active cell proliferation in the stump of joint regeneration (Fig. 4A, B). This result contrasted with the low cell proliferation in the stump region of the lower arm regeneration (Fig. 4B). Our blastema transplantation experiments in frogs indicated that the functionality of joint regeneration was regulated by the stump region (Fig. 5). Abundant joint stump-derived cells contributed to the distal regenerated joint region even after the blastema formation (Fig. 5C-D’). Generally, previous investigations of the stump responses during regeneration focused on the stage until blastema formation (Tsai et al., 2019 and 2020). In limb regeneration, it is believed that the stump makes no response or contribution after blastema formation because of the distance from the regenerative signals such as Fgf from the AEC (Hay and Fischman, 1961; McCusker et al., 2015; Stocum, 2017). However, our results indicate that the stump tissues and/or cells actively contribute to the distal (blastema) region even after blastema formation and this may contribute to the high ability of reintegration in joint regeneration. It has already been indicated that articular cartilage is necessary for regenerating functional joint structure (Tsutsumi et. al., 2015). In that experiment, removing residual articular cartilage after joint cutting. Compared to normal joint regeneration, joint regeneration with articular cartilage removal cannot regenerate functional joints but regenerate complete structures of the distal portion including digits. This result suggested that the proximal portion of the regenerating limb was formed from residual cartilage and its surrounding tissues.

Taken together, our findings suggest that the stump in joint regeneration seemed to behave as a “post-blastema” which is a concept from planarian regeneration (Saló and Baguñà, 1984; Tasaki et al., 2011a and 2011b; Lee et al., 2022). “Post-blastema” designates the proliferating stump tissues after blastema formation, and these tissues can supply cells to the blastema during regeneration. We need to focus more attention on the stump region as an active contribution to the regeneration. Recently, it was shown that the clearance processes of the stump tissues during regeneration are necessary for proper regeneration in the limb (Riquelme-Guzmán et al., 2022). It will be necessary to reveal the role of the stump during the whole regeneration process to understand reintegration during regeneration.

### Signal interactions in joint regeneration

Joint regeneration-specific active responses in the stump region should be driven by some regenerative signal(s). One of the most important signals in regeneration is Fgf (Christen and Slack, 1997; Yokoyama et al., 2000; Cannata et al., 2001; Okumura et al., 2019; Aztekin et al., 2021). Fgf can induce cell proliferation and dedifferentiation in many situations (Maddaluno et al., 2017). Several studies in axolotl indicate that Fgf can regulate dedifferentiation and intercalation (Satoh et al., 2010). Fgf signaling is also related to Mmps expression in the blastema (Satoh et al., 2011). The main source of Fgf is the wound epithelium and the most distal epidermis of the blastema, the AEC (Okumura et al., 2019; Aztekin et al., 2021). Our results indicate that Fgf signaling was received not only in the blastema but also in the stump in joint regeneration (Fig. 1). Furthermore, we also found that Fgf can induce *mmp1* expression in the uninjured articular cartilage (and synovium) (Fig. 6). This indicates that Fgf signaling can induce *mmp1* expression in the articular cartilage of the regenerating joint.

The actual function of ECM degradation in articular cartilage was not clear (Tsutsumi et al., 2015 and 2016). Except for the dedifferentiation, the other possible role of articular cartilage degradation is the release of growth factors from the cartilage ECM. Our study suggests that ECM degradation can be related to Bmp signaling induction (Fig. 7). This Bmp signaling activation may be important for two reasons. The first one is the function of BMPs in blastema formation. Like Fgf signaling, Bmp signaling is essential for limb regeneration (Vincent et al., 2020; Satoh et al., 2022). Moreover, Bmps induce regeneration in co-operation with Fgfs (Makanae et al., 2014). Thus, Bmps secreted from the degraded articular cartilage may possibly function as a blastema-inducing signal and may interact with Fgfs in joint regeneration. The second possible function of Bmps is joint induction in regeneration. Several Bmps can induce joint or joint-like structures in regeneration (Satoh et al., 2005; Yu et al., 2019). It is possible that Bmps secreted from degraded articular cartilage induce joint structures. These signal inductions in the stump and signal interactions between the blastema and stump mediated by tissue degradation and tissue contribution from the stump to the blastema can finally induce reintegration responses of joint regeneration (Fig. 8).

**Fig. 8.**
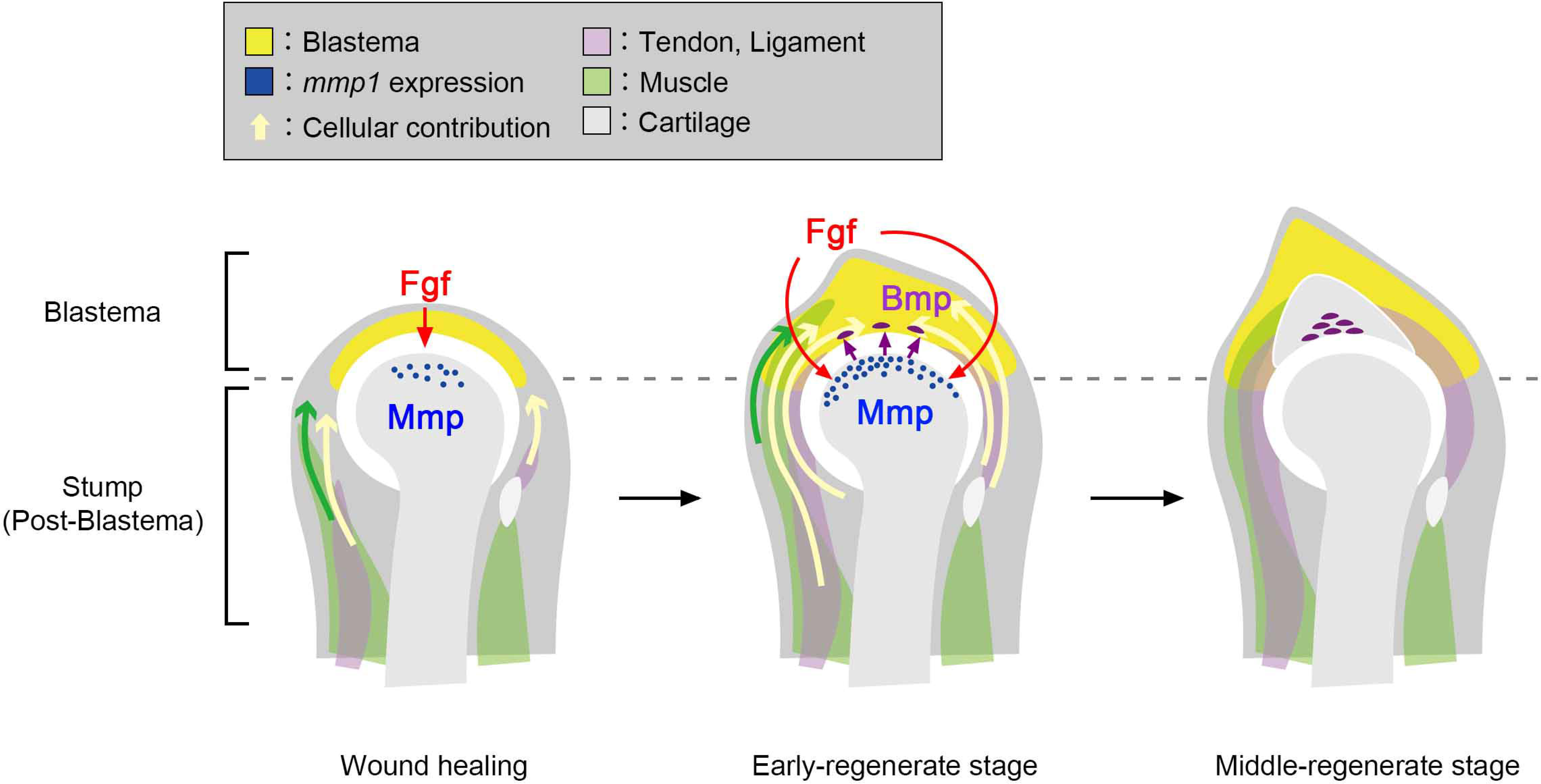
Model of the reintegration mechanism of joint regeneration. In joint regeneration, an active cellular contribution from the stump region derived from several tissues can be seen. Mmp expression in the stump was induced by the regenerative signal Fgf and degraded tissues. Subsequently to the tissue degradation, Bmp signal was upregulated in the boundary between blastema and stump. These reciprocal processes can induce typical reintegration responses in joint regeneration.

### The differences between joint injury and joint regeneration

In humans, it is known that several types of joint injury, including arthritis and rheumatoid arthrosis, occur because of damage and aging (Smolen et al., 2018). Several studies have indicated that partial joint structures show some regenerative activity in mammals, especially in the neonatal period (Yu et al., 2019; Miura et al., 2020). However, the cellular responses and repair of the damage seem to be incomplete in mammals (Caldwell and Wang, 2015). In frogs, joint tissues (articular cartilage, tendon, ligament, and synovium) have higher cellular ability, not only for repair but also dedifferentiation and re-differentiation. Moreover, joint tissues have reactivity to regenerative signals and have the capacity to properly carry out signal cascades in the regenerative situation. Joint regeneration in frogs will thus provide useful information about what results in the difference between functional regeneration and non-functional injury. Recently, in regenerative medicine, organoid formation and organoid transplantation to damaged organs are actively researched (Yui et al., 2012; Fordham et al., 2013; Revah et al., 2022; Westerling-Bui et al., 2022). However, the view of host’s conditions seems insufficient in such studies. It is speculated that a proper connection to the stump and the graft is necessary for functional regeneration even though organoid transplantation. This study is a good model of how to integrate the stump into the newly formed structure. In the future, investigating the mechanism that enables proper, functional regeneration in regenerative animals could provide good material (or strategies) for proper repair and regeneration in regenerative medicine for humans.

## Acknowledgements

This research was supported by JSPS KAKENHI (Grant Number 16H06376 to 21K15145). This research was partly performed at Tottori Bio Frontier managed by Tottori prefecture. We thank the Amphibian Research Center (Hiroshima University) for the kind gift of *Xenopus laevis* adults and Dr. Harumasa Okamoto and Dr. Ikuko Hongo (Gakushuin University) for the kind gift of *Xenopus laevis* embryos. We also thank Dr. Takashi Takeuchi for his generous support and Dr. Gembu Abe for his valuable discussion.

---

**Figure 1 Supplement 1.**
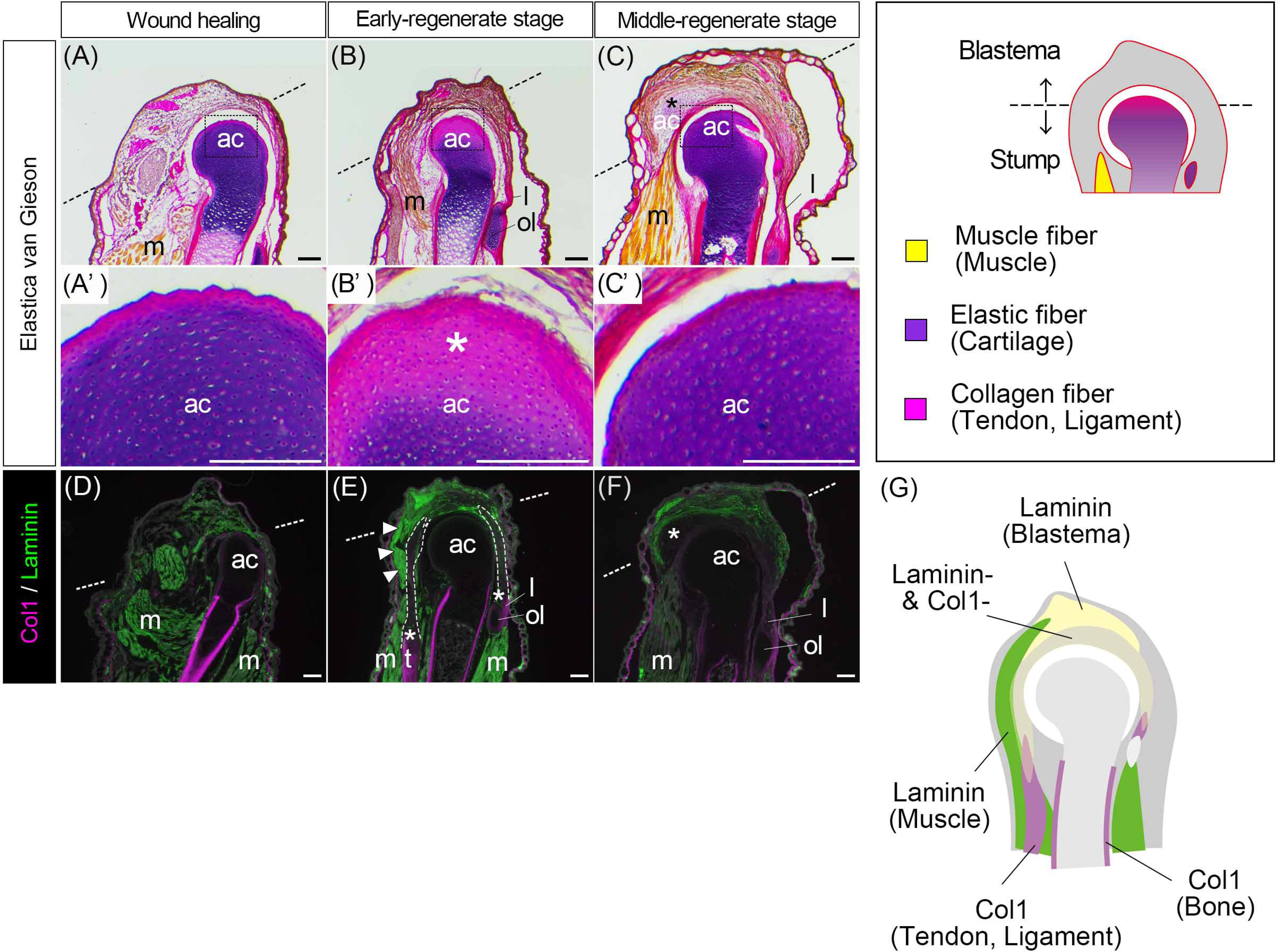
Tissue construction during joint regeneration. Tissue construction changes during the wound healing to the middle-regenerate stage in joint regeneration. (A-C) Joint regeneration staining with EVG. Each color indicates tissue-specific fibers: muscle fiber (yellow), elastic fiber (purple), and collagen fiber (red or magenta). (A’-C’) Elastic fiber conformation in the peripheral region of the articular cartilage of each regenerate in joint regeneration. (B’) The asterisk indicates tissue structure change in articular cartilage. (D-F) Col1 and Laminin expression in joint regeneration. (E) The dotted enclosure indicates proximal to distal contiguous Col1 and Laminin double-negative cells. Asterisks indicate the connected regions of Col1-positive tendon/ligament and Col1/Laminin-double-negative cells. Arrowheads indicate Laminin-positive muscle. (F) The asterisk indicates Col1/Laminin-double-negative cells starting to differentiate cartilage. (G) Schematic diagram of the ECM structure in joint regeneration in the early-regenerate stage. ac, articular cartilage; m, muscle; ol, olecranon; t, tendon; l, ligament. Scale bars, 100 μm.

**Figure 2 Supplement 1.**
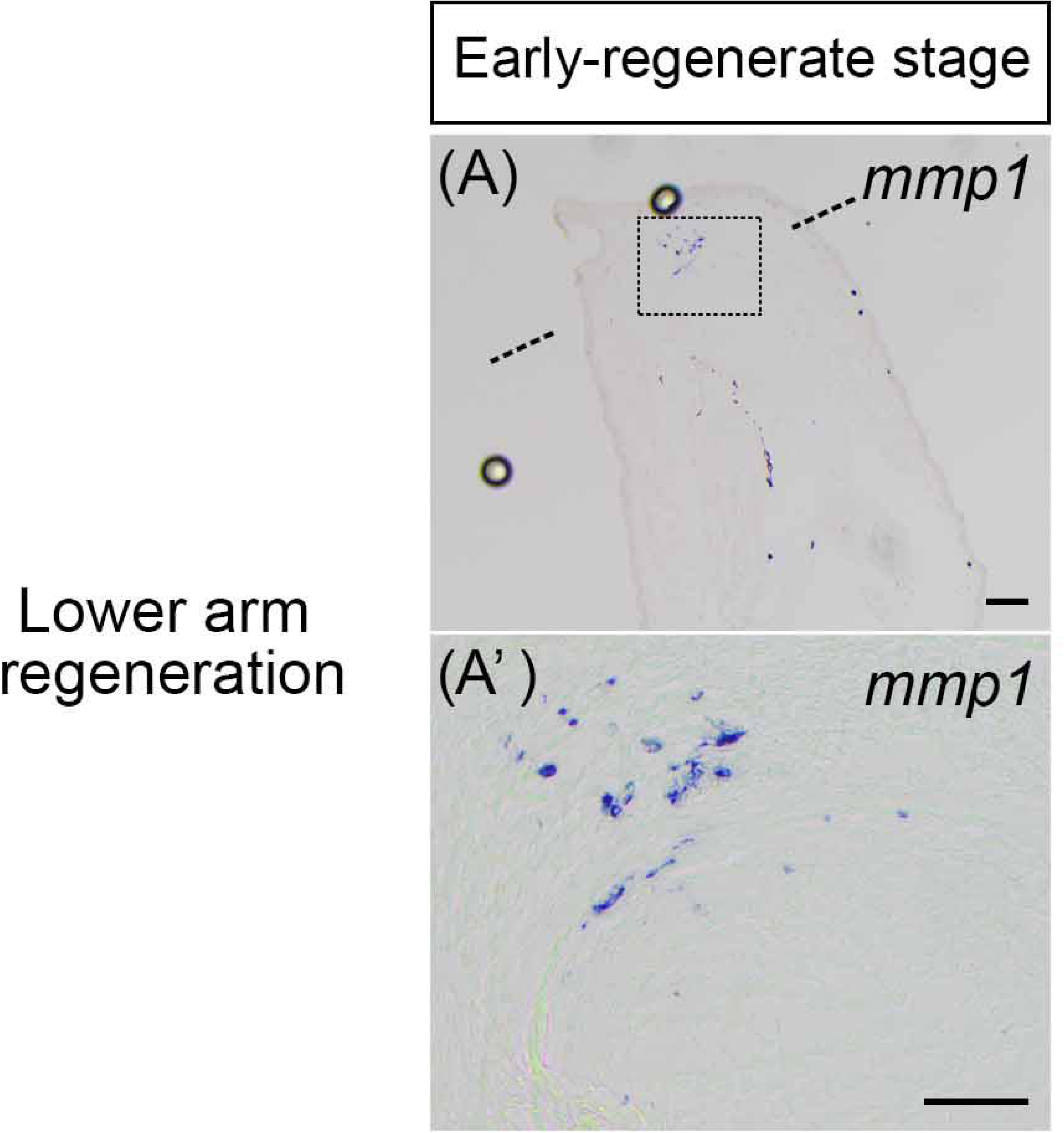
*mmp1* expression of lower arm regeneration. The section *in situ* hybridization to detect *mmp1* expression of the early-regenerate stage in lower arm regeneration. A dashed line indicates a junction of the blastema to the stump. (A’) Highly magnified view of the boxed area in (A). Scale bars, 100 μm.

**Figure 3 Supplement 1.**
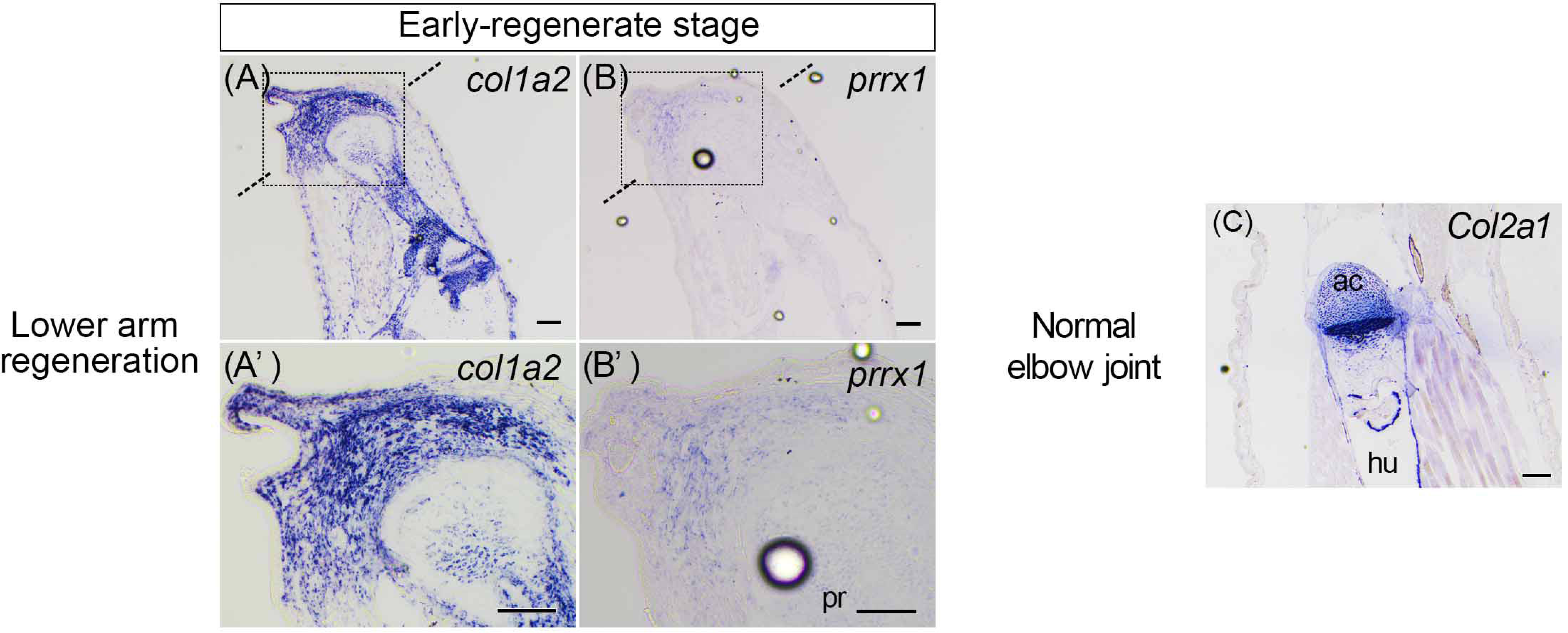
Gene expression patterns of lower arm regeneration and normal elbow joint. (A-B’) The section *in situ* hybridization to detect *col1a2* and *prrx1* expression of the early-regenerate stage in lower arm regeneration. A dashed line indicates a junction of the blastema to the stump. (A’, B’) Highly magnified view of the boxed area in (A, B). (C) *Col2a1* expression in the normal elbow joint. pr, periosteum; ac, articular cartilage; hu, humerus. Scale bars, 100 μm.

